# Commensal-Pathogen Competition Impacts Host Viability

**DOI:** 10.1101/245324

**Authors:** David Fast, Benjamin Kostiuk, Edan Foley, Stefan Pukatzki

**Affiliations:** Department of Medical Microbiology and Immunology, University of Alberta, Edmonton, AB T6G 2S2, Canada.; Department of Immunology & Microbiology, University of Colorado School of Medicine, Aurora, CO 80045.

## Abstract

While the structure and regulatory networks that govern the activity of the type-six secretion system (T6SS) of *Vibrio cholerae* are becoming increasingly clear, we know less about the role of the T6SS in disease. Under laboratory conditions, *V. cholerae* uses the T6SS to outcompete many Gram-negative species, including other *V. cholerae* strains and human commensal bacteria. However, the role of these interactions has not been resolved in an *in-vivo* setting. We used the *Drosophila melanogaster* model of cholera to define the contribution of the T6SS to *V. cholerae* pathogenesis. Here, we demonstrate that interactions between the T6SS and host commensals impact pathogenesis. Inactivation of the T6SS, or removal of commensal bacteria attenuates disease severity. Re-introduction of the Gram-negative commensal bacterium *Acetobacter pasteurianus* into a germ-free host is sufficient to restore T6SS-dependent pathogenesis. Together, our data demonstrate that the T6SS acts on commensal bacteria to promote the pathogenesis of *V. cholerae*.

## Introduction

Diarrheal diseases, including cholera, are the leading cause of morbidity and the second-most common cause of death among children under the age of five worldwide^1,2^. Cholera alone, caused by the marine bacterium *Vibrio cholerae* is responsible for several million cases and over 120,000 deaths annually^3^. When water contaminated with *V. cholerae* is ingested, pathogenic *V. cholerae* bacteria pass through the gastric acid barrier, penetrate the mucin layer of the small intestine, and adhere to the underlying epithelial lining. *V. cholerae* multiplies rapidly, secretes cholera toxin^4^, and exits the human host in immense numbers (˜10^9^ bacteria per mL watery stool)during diarrheal purges^5^.

Despite *V. cholerae*’s numerical inferiority upon arrival in the gut, *V. cholerae* overcomes the barrier presented by commensal gut bacteria, which secrete bacteriocins and compete for nutrients and attachment sites. To establish an infection, *V. cholerae* encodes gene regulation and adaptive responses that permit aggressive expansion in the host. For example, the ToxR regulon acts in concert with other signaling cascades to subvert host homeostasis^6^.

To maintain competitiveness in microbial communities, *V. cholerae* uses its type six secretion system (T6SS), a molecular system that delivers toxic effectors into prokaryotic and eukaryotic prey. If the target cell lacks cognate immunity proteins, it rapidly succumbs to the injected toxin, allowing *V. cholerae* to dominate a niche^7–9^. The T6SS selectively targets Gram-negative bacteria and eukaryotic phagocytes such as macrophages, providing *V. cholerae* a competitive advantage within the small intestine^10–12^. In contrast, Gram-positive bacteria are immune to T6SS-mediated toxicity, potentially due to their thick, protective peptidoglycan layer^13,14^. Studies with other bacteria suggest that pathogens deploy their T6SS to overcome the barrier presented by commensal bacteria^15^. For example, *Salmonella enterica Serovar Typhimurium* (*S. typhimurium*) uses its T6SS to outcompete Gram-negative commensals and enhances colonization of the adult mouse gut^15^. Alternatively, the *Campylobacter jejuni* T6SS is thought to act directly on eukaryotic cells to support persistent *in-vivo* colonization of IL-10 deficient mice^16^.

Studies with the infant mouse and rabbit models showed that the T6SS of *V. cholerae* is functionally active inside the host^17–19^, and that the T6SS of *V. cholerae* contributes to inflammation in the infant mouse model^20^. Furthermore, gene expression profile data from humans infected with V. cholerae showed an upregulation of T6SS genes^21^. Despite the experimental support for T6SS activation inside the host, we do not know if the T6SS acts on commensal bacteria during an infection, or how relevant such interactions are for disease progression. To address this question, we used the model organism, Drosophila melanogaster, to determine the influence of the T6SS on the virulence of *V. cholerae*, and to examine interactions between commensal bacteria and *V. cholerae* during an enteric infection. The *Drosophila-Vibrio* model is particularly suited for such studies. Flies naturally die from *Vibrio* infection; the gut microbiome of flies is simple, and easily manipulated. Importantly, fly gut physiology is similar to more complex vertebrate systems^22–31^.

We found that the T6SS-positive El Tor strain C6706 establishes a lethal cholera-like disease in adult flies. Inactivation of T6SS activity significantly impaired host colonization, reduced disease symptoms, and extended host survival. T6SS-dependent killing of *Drosophila* specifically requires *Drosophila* association with the Gram-negative commensal bacterium, *Acetobacter pasteurianus* (*Ap*). Removal of commensal bacteria abrogates T6SS-mediated killing of the host, and reintroduction of *Ap*, either alone, or in combination with addition commensals, fully restores T6SS-dependent lethality. Collectively, our work establishes that interactions between the T6SS and commensal Gram-negative bacteria contributes to the progression of disease in *Drosophila*.

## RESULTS

### The T6SS interacts with commensal bacteria to influence host viability

As *Drosophila* is naturally susceptible to infection with *V. cholerae*, we reasoned that the fly model provides a platform to determine the function of the T6SS function *in vivo*^22^. Pandemic *V. cholerae* strains belong to two biotypes of the O1 serogroup. The classical biotype responsible for the first six pandemics carries multiple nonsense mutations and deletions in the T6SS genes, resulting in a disabled T6SS (D.U., B.K. and S.P., unpublished observation). The El Tor biotype responsible for the current seventh cholera pandemic has a functional T6SS that becomes activated upon host entry. Thus, we tracked the survival of adult flies that we infected orally with El Tor strain C6706, or strain O395. Both biotypes encode acetyl-CoA synthase, and cholera toxin, bacterial products that contribute to pathogenesis in *Drosophila*^22,32^. C6706 has been shown to be avirulent in the fly model due to the activity of the regulator HapR^33^. However, our laboratory isolate of C6706 effectively kills adult *Drosophila* due to decreased *hapR* levels. As controls, we measured the viability of adult flies raised on sterile, bacterial growth medium. Under these conditions, infection with O395 caused a moderate reduction in adult viability compared to controls (Fig. 1a). In contrast, C6706 had a significant impact on the lifespan of infected adults. The median viability of C6706-infected flies was a third of that observed for controls (50hrs vs 149hrs, Fig. 1a), and all C6706-infected flies perished within seventy-two hours of infection (compared to 170 hours for mock-infected flies).

**Figure 1.**
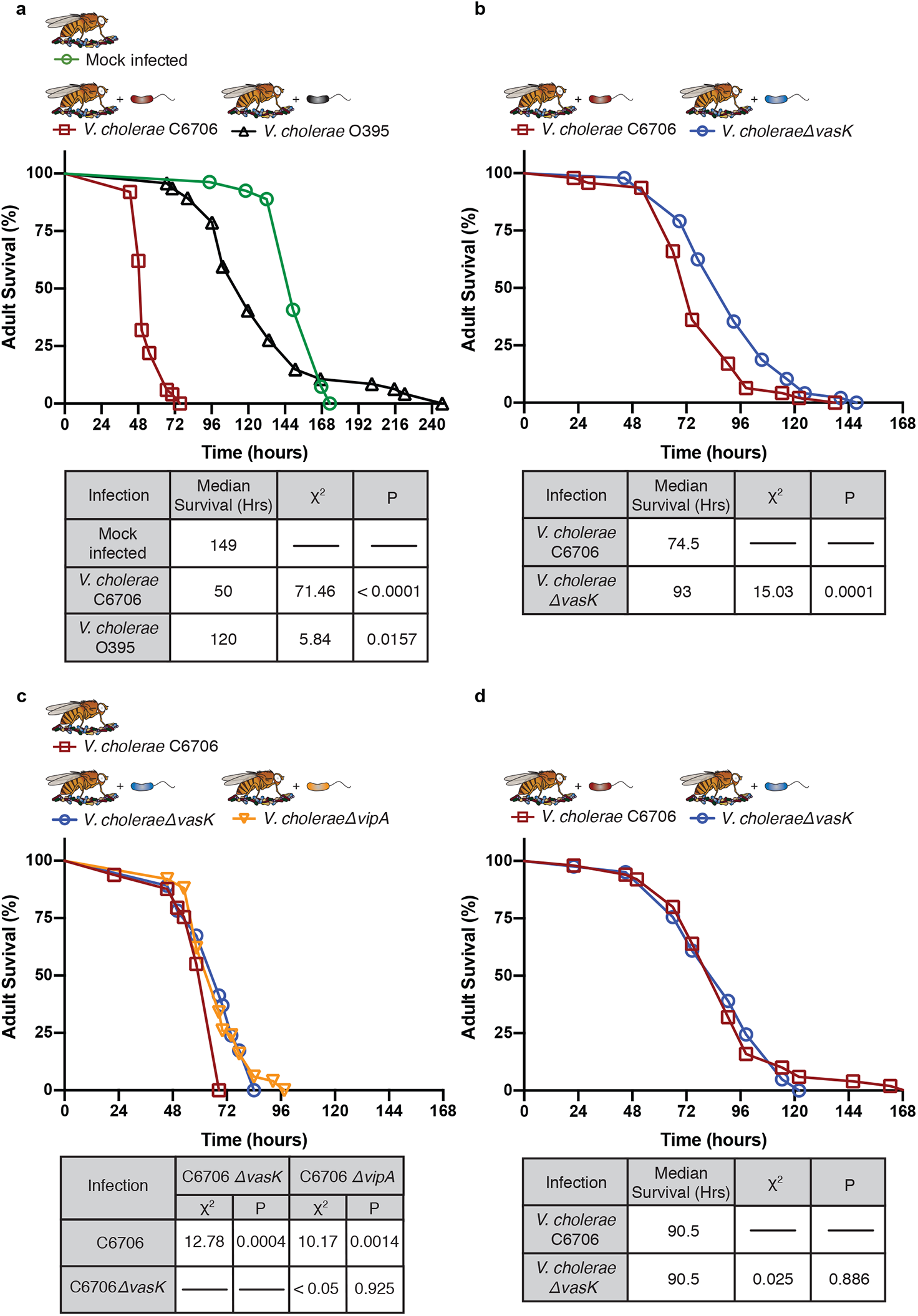
The T6SS contributes to the pathogenesis of *V. cholerae* in a commensal-dependent manner. **a**, Survival curve of female CR *w^1118^* flies raised at 29°C for 5–6 days, and infected with strains of *V. cholerae* responsible for two distinct pandemics. LB alone served as mock infection. **b**, Survival curve of adult conventionally reared (CR) flies infected with T6SS functional (C6706) or T6SS nonfunctional mutant (C6706ΔvasK) *V. cholerae*. **c**, Survival curve of adult CR flies infected with T6SS functional (C6706) and two nonfunctional T6SS mutants (C6706ΔvasK and C6707ΔvipA) **d**, Survival curve of adult germ free (GF) flies infected with functional T6SS (C6706) or nonfunctional T6SS mutant (C6706*ΔvasK*) of *V. cholerae*. **d**, was same performed at the same time and infected with the same bacterial cultures as b. For each graph, the y-axis represents percent survival and the x-axis represents infection time in hours. Tables are results of Long-rank (Mantel-Cox) test of graph data. **a**, χ2 and P values are relative to mock infected flies **b-d**, χ^2^ and P values are relative to wild-type C6706 infected flies. n = 50 per group, for all experiments.

As C6706 encodes a functional T6SS, we asked if disabling the T6SS affects *V. cholerae* pathogenesis. We infected adult flies with wild-type C6706, or with C6706 carrying an in-frame deletion of *vasK*, which encodes an inner membrane protein essential for T6SS assembly^13^. We found that disabling the T6SS in C6706 significantly impaired pathogenesis (Fig. 1b). Deletion of *vipA*, which encodes a protein that makes up the outer sheath of the T6SS infection machine^34^, also impaired pathogenesis (Fig. 1c). Both T6SS mutations had near-identical attenuating effects on host killing. Combined, these results establish that the T6SS contributes to *V. cholerae* pathogenesis *in vivo*. Inactivation of the T6SS does not abolish pathogenesis, suggesting that *V. cholerae* employs additional virulence factors such as cholera toxin, metabolite production, and quorum sensing^22,32,33^ to kill the host in a T6SS-independent manner.

As the T6SS targets eukaryotic and prokaryotic cells^10^, we asked if the T6SS contributes to host killing either by direct effects on the host, or by indirect effects on the intestinal microbiota^10,13,20^. We examined survival rates of conventionally-reared (CR) and germ-free (GF) flies that we challenged with C6706 or C6706Δ*vasK*. If the T6SS acts directly on the fly, we expect that removal of commensal bacteria will not affect T6SS-dependent killing of the host. Instead, we found that an absence of commensal bacteria impaired C6706-dependent killing to the point that it was no longer distinguishable from C6706Δ*vasK* (Fig. 1d), indicting that T6SS-dependent killing of a fly host requires the presence of commensal bacteria.

### The T6SS contributes to disease symptoms

Loss of the T6SS impairs *V. cholerae* pathogenesis. We therefore monitored how the T6SS impacts the development of watery, pathogen-laden diarrhea - the hallmark symptom of cholera. We supplemented the infection culture of *V. cholerae* with a nontoxic, non-absorbable, blue dye. We then infected flies for twenty-four hours, and monitored defecation hourly for the next four hours^35^. We first counted individual blue dots to determine defecation frequency. As controls, we measured defecation by uninfected flies that we raised on a solid fly culture medium with blue dye, or on bacterial growth medium supplemented with the same dye. We observed no difference in defecation frequency between flies raised on solid or liquid diets, confirming that the bacterial growth medium alone does not cause diarrhea (Fig. 2a). Likewise, O395 had no measurable effects on defecation frequency (Fig. 2a). In contrast, we found that C6706 caused a drastic increase in the number of fecal marks per fly (Fig. 2a). Similarly, we found an increase in the number of fecal marks per fly from flies infected C6706Δ*vasK*. However, this increase was less pronounced than that of flies infected with C6706. To better assess the contributions of the T6SS to defecation frequency, we performed a linear regression analysis on each of the groups indicated in Fig. 2a. Here, we noticed a significant increase in the number of fecal marks/fly over time from flies infected with C6706, but not from mock-infected flies. Furthermore, there was a significantly lower increase in the number of fecal marks/fly from C6706*ΔvasK*-infected flies, and a smaller portion of these fecal marks could be attributed to infection with C6706Δ*vasK* than C6706 (Fig. 2a). However, T6SS inactivation does not abate diarrheal symptoms likely due to *V. cholerae’s* ability to produce cholera toxin. Together, these data indicate that the T6SS impacts the severity of diarrheal symptoms in infected flies.

**Figure 2.**
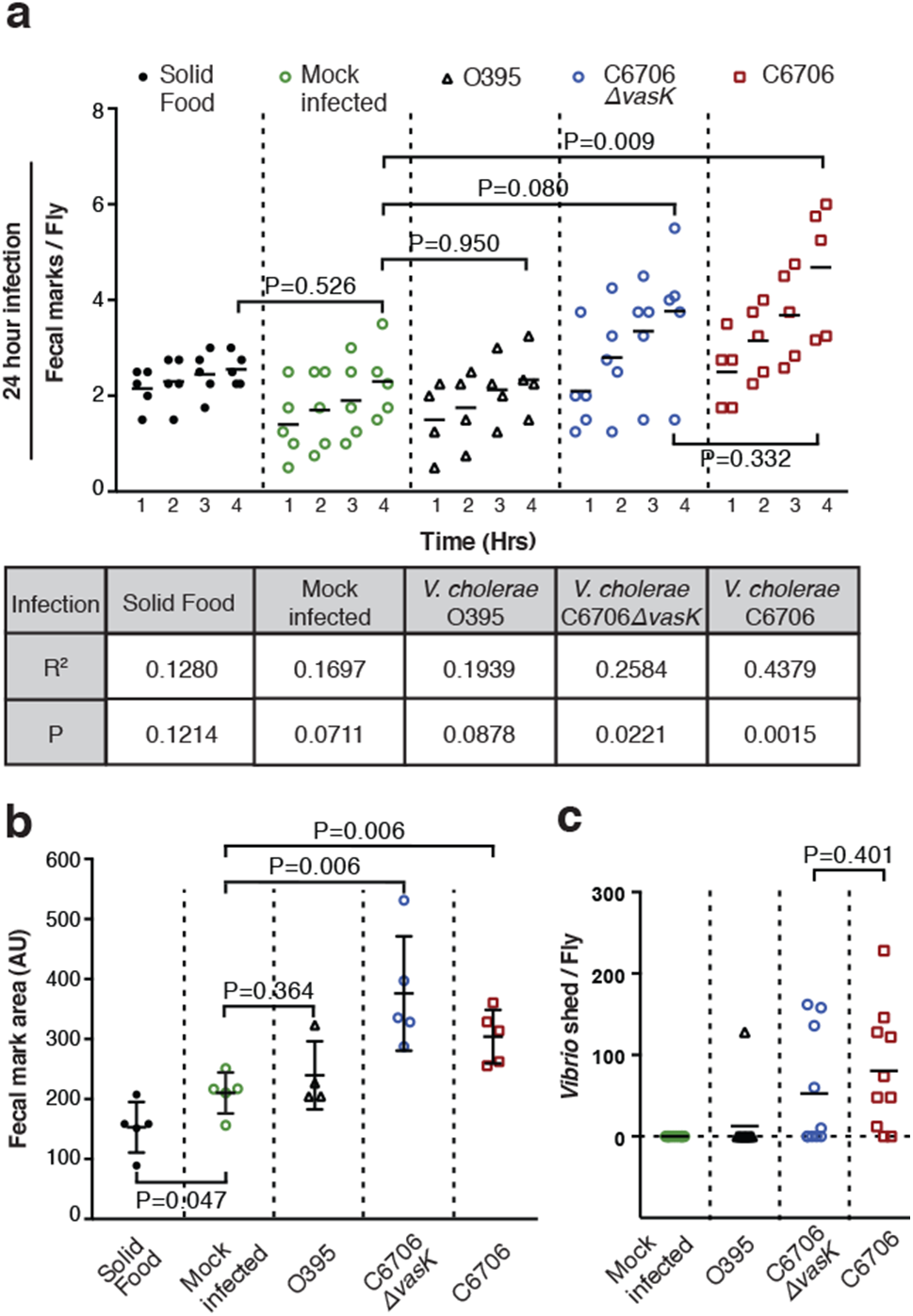
The T6SS contributes to cholera-like disease. **a**,Fecal marks were scored for *w^1118^* flies raised at 29°C for 5–6 days flies fed solid fly food, LB broth, or infected with O395, C6706*ΔvasK*, or C6706 for 24 hours. For **b**, the table below is the result of a linear regression analysis of each treatment group and P values on the graph are the result of a Student’s t-test at 4 hours. **c**, Fecal mark area in μm of spots counted. Each point is the average area taken from a given replicate. Statistics are the results of a Student’s t-test of each group compared to solid food. **d**, *Vibrio cholerae* shed per fly of 5–6 day old *w^1118^* flies fed LB or infected with *V. cholerae* C6706, C6706*ΔvasK*, and O395 for 24 hours. Each data point is the number of *Vibrio* isolated from fecal matter of a single fly.

We also measured the T6SS effects on defecation volume. For this assay, we calculated the surface area of each dot as a proxy for water content of feces. We observed an increase in the area of fecal spots from mock-infected flies raised on a liquid diet compared to flies raised on a solid diet (Fig. 2b). Furthermore, O395 did not cause a significant increase in fecal volume. In contrast, both C6706 and C6706*ΔvasK* significantly increased fecal volume relative to mock infected controls (Fig. 2b), confirming enhanced diarrheal disease in flies infected with either strain. Finally, as the shedding of *V. cholerae* in fecal matter accompanies diarrhea, we quantified the number of *V. cholerae* bacteria excreted by flies that we challenged with the different strains of *V. cholerae*^4^. Whereas we only detected V. cholerae in the feces of a single fly infected with the O395 strain, we found that 8/10 flies infected with C6706 shed *V. cholerae*. Consistent with contributions of the T6SS to disease severity, we only found 5/10 C6706*ΔvasK* infected flies shed the bacteria. In short, our results establish a role for the T6SS in diarrheal symptoms during a *V. cholerae* infection – loss of the T6SS causes a reduction in defecation frequency, and less shedding of *V. cholerae* in the feces of infected animals.

### T6SS promotes intestinal epithelial damage

During infection with *V. cholerae*, diarrhea is accompanied by ultrastructural changes to the host intestinal epithelium^36^. Therefore, transmission electron microscopy was used to examine the posterior midgut (analogous to the small intestine of mammals) ultrastructure of mock-infected flies, or flies challenged with C6706 or C6706Δ*vasK* for fifty hours. Mock-infected intestines had a readily identifiable lumen, an epithelium of evenly-spaced columnar cells with extensive brush borders, and morphologically normal nuclei and mitochondria (Fig. 3a-f). In contrast, an intact intestine could no longer be discerned in flies challenged with C6706 (Fig. 3g-i). Instead, the gut consisted of a disorganized mass of host cells that lacked apical brush borders, and completely engulfed the intestinal lumen. Extensive shedding of epithelial structures into the presumptive lumen (boxes, Fig. 3g-i, h-j) was observed, and high magnification images revealed characteristics of cell death, such as nuclear decondensation, and swollen mitochondria (Fig. 3k, l). Infection with C6706Δ*vasK* caused an intestinal phenotype that was intermediate between mock-infected controls and C6706-infected adults. Guts infected with C6706Δ*vasK* retained elements of intestinal organization, such as identifiable epithelial cells with brush borders, and a recognizable lumen (Fig. 3m-p). Furthermore, challenges with C6706Δ*vasK* did not have obvious impacts on nuclear or mitochondrial organization (Fig. 3q, r). However, we did notice intestinal phenotypes similar to flies that were challenged with C6706. Specifically, infection with C6706Δ*vasK* caused an extrusion of epithelial cell matter into the lumen (dashed boxes, Fig. 3m, n), a phenotype consistent with pathogen-mediated destruction of host epithelial structures in *Drosophila*^37^. In summary, these results uncover a role for the T6SS in the severity of disease in adult *Drosophila*. Inactivation of the T6SS diminishes damage to the intestinal epithelium, lowers the severity of diarrhea, and extends host mortality times.

**Figure 3.**
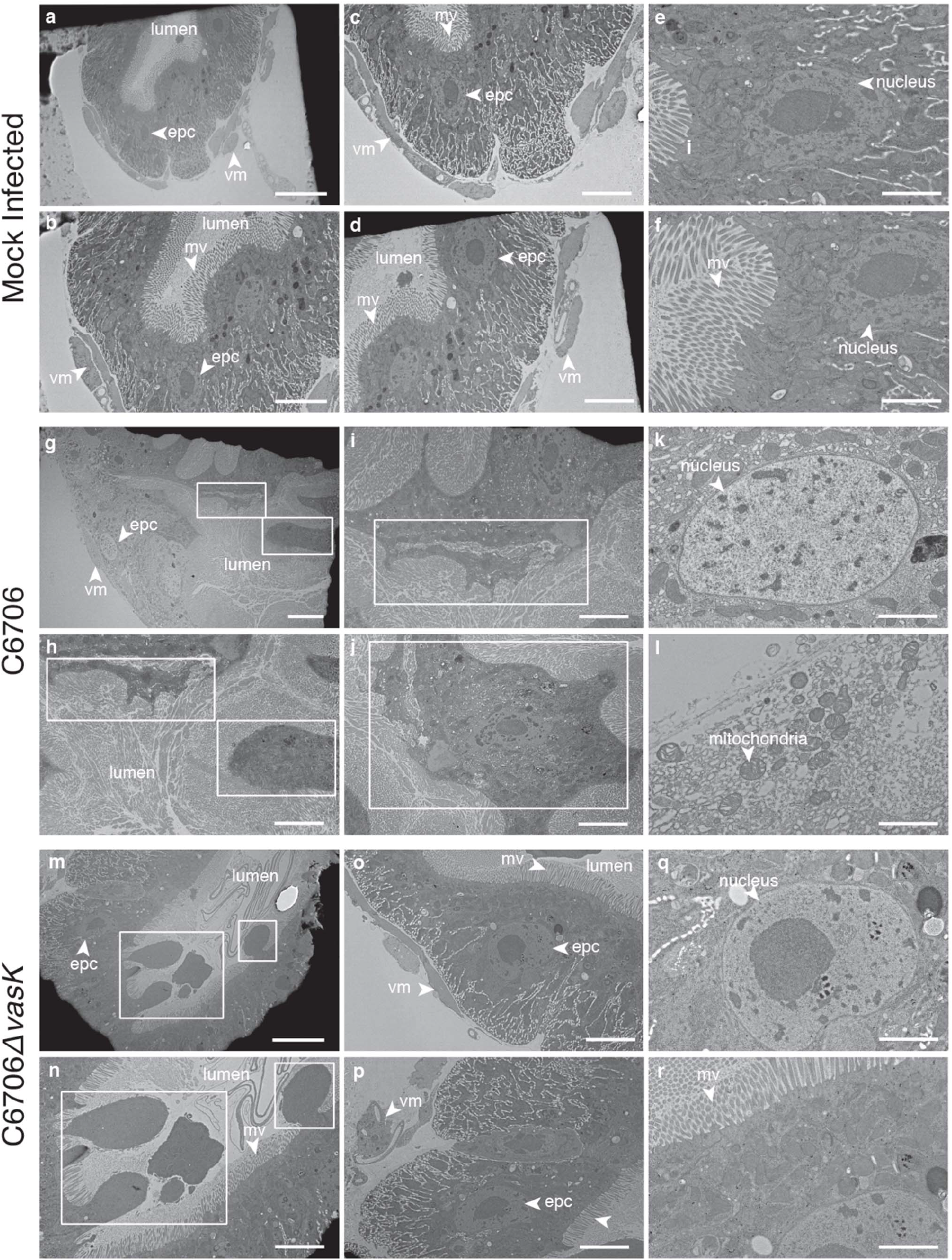
T6SS contributes to *V. cholerae* intestinal pathogenesis. Transmission electron microcopy of the posterior midguts of flies, mock infected (first and second row) or infected with C6706 (third and fourth row), C6706*ΔvasK* (fifth and sixth) after 50 hours of infection. Large scale bars are 10μm small scale bars are 5μm. The microvilli are marked mv, the epithelial cells epc, and the visceral muscle vm. Cells protruding into the lumen are indicated with a dashed line box.

### The T6SS influences pathogen-commensal interactions in the intestine

As T6SS-assisted pathogenesis requires an intact microbiota (Fig. 1d), we asked if intestinal bacteria influence host colonization by *V. cholerae*. Laboratory-reared *Drosophila* typically have low diversity gut bacterial consortia^23–25,38^. In our laboratory, wild-type fly intestines are dominated by the Gram-negative commensal *Acetobacter pasteurianus* (*Ap*), and the Gram-positive *Lactobacilli, L. brevis* (*Lb*) and *L. plantarum* (*Lp*)^39^. To determine if *Ap* or *Lactobacilli* influence host colonization by *V. cholerae*, we established populations of germ-free adult flies, and adults that we associated exclusively with *Ap* or *Lb* (Fig. 4a). We then challenged the respective populations with C6706 or C6706*ΔvasK*, and measured the colony forming units per fly (CFU/Fly) of *V. cholerae* as a function of time. We found that C6706 and C6706*ΔvasK* were equally effective at colonizing GF intestines, or intestines that exclusively contained *Lb* (Fig. 4b, c). In each case, the numbers of C6706 and *C6706ΔvasK* increased over time and reached nearly identical levels at 24 hours of infection (Fig. 4b). These data indicate that the T6SS is dispensable for the colonization of a germ-free gut, or a gut that houses the Gram-positive bacteria *Lb*. In contrast, removal of the T6SS significantly impaired the ability of *V. cholerae* to colonize an adult intestine that we pre-associated with the Gram-negative commensal, *Ap*. In this scenario, C6706 titres increased significantly from 6 to 24 hours of infection in the intestines of *Ap*-colonized adults. In contrast, there was no increase in the load of C6706*ΔvasK* from 6 to 24 hours (Fig. 4d). By 24 hours, we found an appreciable, though not statically significant, difference in CFU/Fly between C6706 and C6706*ΔvasK* (Fig. 4d). These data indicate that the T6SS supports colonization of intestines that exclusively contain *Ap*.

**Figure 4.**
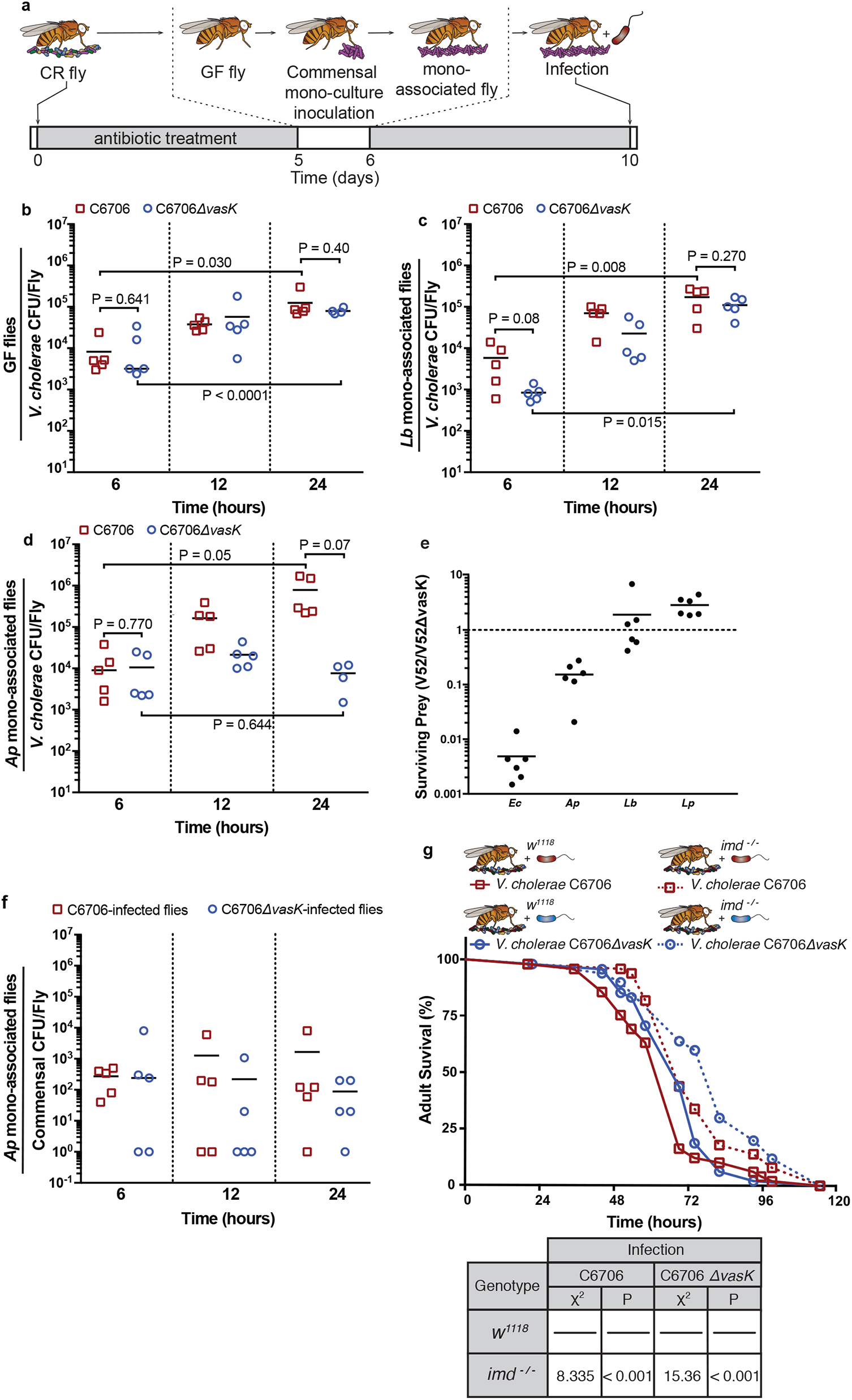
Composition of the microbiome determines T6SS-mediated gut infection. **a**, Schematic representation for the generation of adult mono-associated flies. Colony forming units per fly of *V. cholerae* strains C6706, C6706*ΔvasK* of surface-sterilized **b**, GF flies, **c**, *Lb* mono-associated flies, **d**, *Ap* mono-associated flies at indicated time points. Each data point represents a biological replicate of 5 flies. P values are the result of a Student’s t-test. **e**, An *in-vitro* competitive assay between *V. cholerae* V52 and V52Δ*vasK* against *E. coli* as a positive control and commensal bacteria *Ap, Lb* and *Lp*. Bacteria were co-incubated for 4 hours at 37C. Surviving prey bacteria in the presence of the T6SS was divided by the surviving prey in the absence of the T6SS (Δ*vasK*). **f**, Colony forming units per fly of *Ap* from flies infected with C6706 or C6706*ΔvasK*. Each data point represents a biological replicate of 5 flies. **g**, Survival curve of female CR w1118 or *imd^-/-^* flies raised at 29°C for 5–6 days, and infected with C6706 or C6706Δ*vasK*. Tables are results of Long-rank (Mantel-Cox) test of graph data, χ^2^ and P values are relative to *w^1118^* infected flies.

As the T6SS assists *V. cholerae* colonization of a gut associated with *Ap*, we asked if the T6SS kills *Ap* in in a standard laboratory killing assay^13^. As shown in Fig. 4e, *V. cholerae* effectively killed the T6SS-susceptible prey *Escherichia coli* K12 strain MG1655 in this assay as shown previously^13^. Furthermore, we saw no evidence of T6SS-dependent killing of either *Lp*, or *Lb* (Fig. 4e). This is consistent with previous observations that Gram-positive bacteria are naturally refractory to T6SS activity^13,14^. In contrast, we noticed substantial T6SS-dependent killing of *Ap* by *V. cholerae* (Fig. 4e). These data raise the possibility that the T6SS facilitates host colonization through eradication of commensal *Ap*. To test this hypothesis, we measured total *Ap* numbers in the intestines of flies that we mono-associated with *Ap* and then challenged with C6706 or C6706*ΔvasK*. We did not detect obvious impacts on *Ap* numbers in the presence of T6SS-positive C6706, suggesting that *V. cholerae* infection does not have a substantial effect on total commensal numbers (Fig. 4f).

As we detected no change in *Ap* numbers, we tested the alternative hypothesis that T6SS-mediated killing of a subset of intestinal *Ap* engages secondary responses in the host that accelerate death. For example, we (Fig. 4g) and others found that mutations in the Immune Deficiency (IMD) antibacterial pathway attenuate *V. cholerae*-dependent killing^40^. To determine if T6SS-mediated killing of the host involves pathological activation of immune responses, we infected wild-type, and *imd* mutant flies with C6706, and C6706Δ*vasK*. Mutation of either *vasK* or *imd* prolonged host viability to near equal extents (Fig. 4g). Ablation of the T6SS in combination with an *imd* mutation extended host viability further (Figure 4g). Together, these data suggest that additive effects from the T6SS of *V. cholerae* and the IMD pathway of *Drosophila* synergistically control host viability.

### The microbiome directly influences T6SS-dependent pathogenesis

The T6SS contributes to *Drosophila* killing by *V. cholerae* (Fig. 1b,c), T6SS-assisted killing of *Drosophila* requires an intestinal microbiome (Fig. 1d) and the T6SS specifically targets the Gram-negative commensal *Ap* (Fig. 4e). These observations led us to ask if interactions between the T6SS and *Ap* are a prerequisite for T6SS-mediated killing of the host. To test this hypothesis, we examined host viability in populations of adult flies that we associated exclusively with *Ap*, or *Lb*, and subsequently infected with C6706 or C6706Δ*vasK*. For each study, we ran a parallel infection study on CR flies with the same cultures of *V. cholerae*. Loss of the T6SS significantly impaired pathogenesis in each test with control, CR flies (Fig. 5d, e, i). However, loss of the T6SS did not diminish *Vibrio* pathogenesis in adult flies that we associated exclusively with *Lb* (Fig. 5a). As *Lb* also fails to block host colonization by a T6SS-defective C6706 strain (Fig. 4c), our data suggest that interactions between the T6SS and *Lb* have minimal relevance for host viability. In contrast, we detected significant involvement of the T6SS in the extermination of adults that we mono-associated with *Ap* (Fig. 5b), indicating that *Ap* is sufficient for T6SS-mediated killing of the host. We then asked if Gram-positive commensals can protect *Drosophila* from T6SS-dependent killing of *Ap-*associated flies. Here, we associated adult *Drosophila* with a 1:1:1 mixture of *Ap, Lb*, and *Lp*. We then challenged the poly-associated flies with C6706 or C6706Δ*vasK*, and measured survival rates. In this experimental set, we found that the addition of Gram-positive commensals does not impact T6SS-dependent killing of the host, suggesting that the presence of the common fly commensal *Ap* renders *Drosophila* sensitive to T6SS-dependent killing of the host irrespective of the presence of additional commensals.

**Figure 5.**
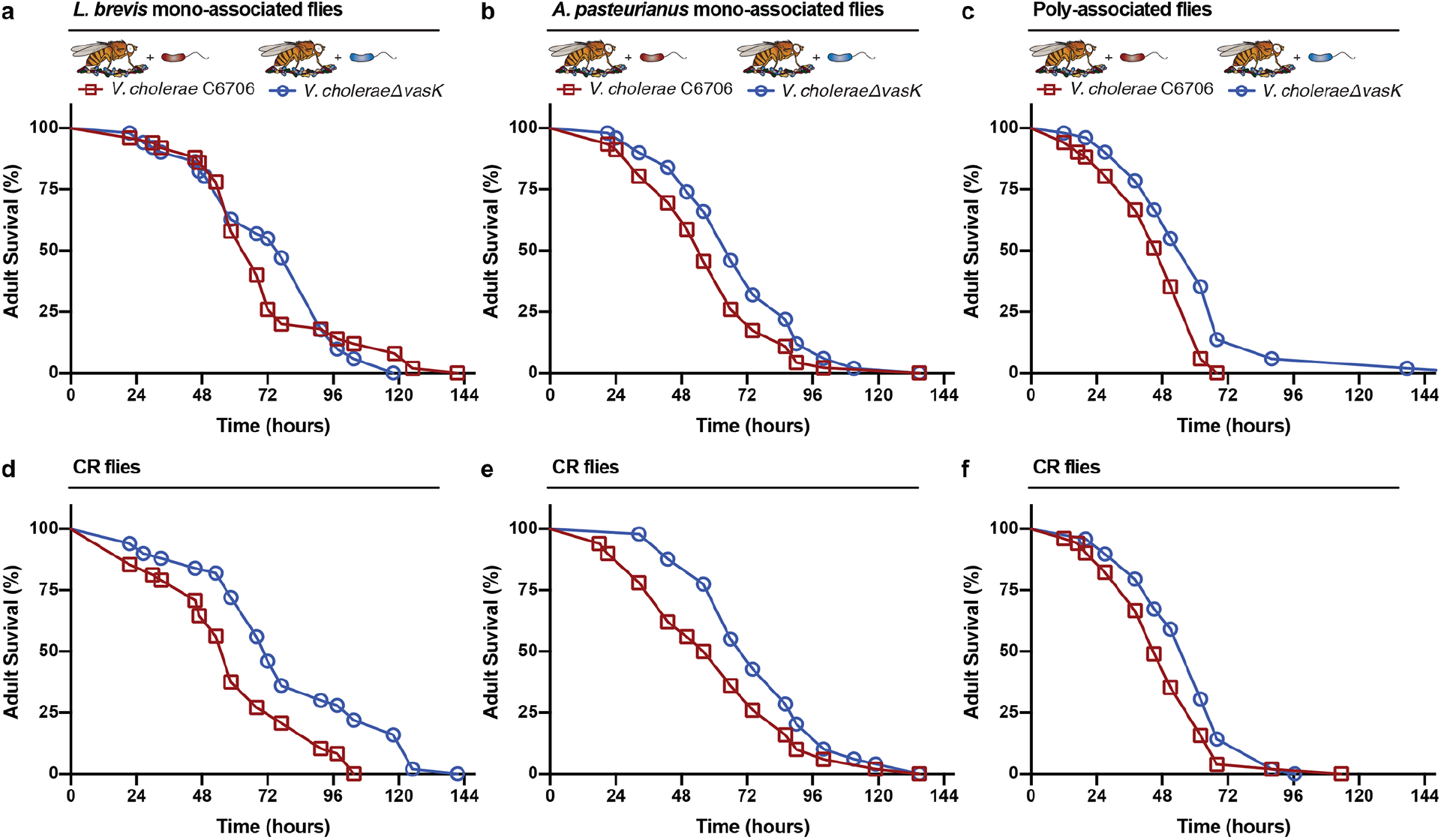
Composition of commensal microbes impacts T6SS virulence contributions *in-vivo*. **a**, survival curves for adult flies mono-associated with *Lb* and infected with C6706 or C6706Δ*vasK*. **b**, survival curves for adult flies mono-associated with *Ap* and infected with C6706 or C6706Δ*vasK*. **c**. survival curves for adult flies poly-associated with *Lb, Ap*, and *Lp* and infected with C6706 or C6706Δ*vasK*. **d-f** and are survival curves of the experiments respective conventionally reared flies controls. For each graph, the y-axis represents percent survival and the x-axis represents infection time in hours. Tables are results of Long-rank (Mantel-Cox) test of graph data.

### Discussion

A dense community of commensal bacteria direct growth, metabolism, and immune activity in the host intestine^41–43^. Under homeostatic conditions, this bacterial community also forms a protective barrier that shields the host from microbial invaders. Thus, pathogenic microbes must navigate a complex preestablished microbiota to establish a productive infection. Here, we used *Drosophila melanogaster* as a simple model to address how pathogen-commensal interactions contribute to host death. We were particularly interested in the T6SS of *V. cholerae*, a molecular syringe with the potential to neutralize host cells or Gram-negative commensals during an infection^13,17^. Using this model, we found that the T6SS contributes to *V. cholerae* pathogenesis, and that the T6SS accelerates host death by interactions with the commensal microbe *Acetobacter pasteurianus*. Removal of either the T6SS, or *Ap* from the *Vibrio-Drosophila* model extends the viability of infected adults, and inoculation of adult flies with *Ap* is sufficient to restore T6SS-dependent killing of the host. These results demonstrate an *in-vivo* contribution of the T6SS to *V. cholerae* pathogenesis.

We found that removal of all intestinal bacteria does not improve host killing by T6SS-deficient *V. cholerae*, arguing against a simple replacement reaction where *V. cholerae* expands into a vacant niche left behind after T6SS-mediated killing of *Ap*. Instead, we found that removal of commensal bacteria attenuated host killing by wild-type *V. cholerae*, suggesting that the presence of commensal bacteria is essential for T6SS-dependent killing of the host. Furthermore, we did not see a substantial drop in *Ap* titres in flies challenged with *V. cholerae*, indicating that *Ap* persists during infection. Finally, we found that inoculation of GF adults with *Ap*, either alone or in combination with *Lactobacilli*, was sufficient to restore T6SS-dependent killing of the host. These observations are in line with a model where T6SS-mediated killing of intestinal *Ap* initiates secondary events that enhance host destruction by *V. cholerae*. We consider the host Immune Deficiency (IMD) pathway a possible mediator of such an effect. IMD is highly similar to the mammalian TNF pathway, and IMD regulates intestinal immune responses to Gram-negative bacteria^44–46^. We showed that loss-of-function mutations in the IMD pathway extend the lifespans of flies infected with wild-type *V. cholerae* (C6706) to a similar extent as wild-type flies infected with a T6SS-negative strain (C6706*ΔvasK*). *Imd* mutant flies infected with the same T6SS-negative strain (C6706) lived even longer, suggesting that activation of IMD and T6SS-dependent killing of *Ap* programs the intestinal environment in a manner that supports *V. cholerae* pathogenesis. The system presented in this report presents an ideal option to characterize T6SS-commensal interactions in a simple host.

## MATERIALS AND METHODS

### Bacterial Strains

All *Drosophila* commensal bacteria strains used were isolated from wild-type lab flies from the Foley lab at the University of Alberta and are as follows: *Lactobacillus plantarum* KP (DDBJ/EMBL/GenBank chromosome 1 accession CP013749 and plasmids 1–3 for accession numbers CP013750, CP013751, and CP013752, respectively), *Lactobacillus brevis* EF (DDBJ/EMBL/GeneBank accession LPXV00000000), and *Acetobacter pasteurianus* AD (DDBJ/EMBL/GeneBank accession LPWU00000000) and are described in^47^. *Lactobacillus* strains were grown in MRS broth (Sigma Lot: BCBS2861V) at 29°C and *Acetobacter pasteurianus* was grown in Mannitol broth (2.5% n-mannitol. 0.5% yeast extract, 0.3% peptone) 29°C with shaking. *Vibrio cholerae* strains were grown in Lysis Broth (LB) broth (1% tryptone, 0.5% yeast extract, 0.5% NaCl) at 37°C with shaking. As indicated, bacteria were grown in the presence of 100 μg/ml streptomycin.

### Oral Infection with *V. cholerae*

Virgin female flies were separated from male flies after eclosion and placed on autoclaved standard Bloomington food for 5 days at 29°C. Flies were starved 2 hours prior to infection. Ten flies were then placed in a vial containing a cotton plug soaked with *V. cholerae* (OD_600_ of 0.125) in groups of 5 per experimental group, resulting in 50 flies for each survival curve. Dead flies were counted every 8hours for the first 100 hours, and every 24 hours thereafter.

### Fly Fecal Assays

Flies were infected as described above except that the infection culture contained Erioglaucine disodium salt (0.1%). After 24 hours of infection, flies were placed in a petri dish lined with a filter paper. The number of new blue spots were counted every hour. After 4 hours, the filter paper was imaged and the size of the spots was calculated with CellProfiler (3.0.0, Broad Institute).

### *V. cholerae* shedding assay

Flies were infected as above. After a 24-hour infection, individual flies were placed in a 96 well plate where each well had been lined with filter paper soaked in PBS + 5% sucrose. After 4 hours, the filter paper was vortexed vigorously for 15 seconds in 1mL of LB and serial dilutions were made on LB + Streptomycin. CFUs were counted the next day.

### Fly husbandry

All experiments were performed with virgin female flies. *w^1118^* flies were used as wild type. The *imd^-/-^* (*imd[*Ey08573) flies have previously been described^48^. Flies were raised on standard corn meal medium (Nutri-Fly Bloomington Formulation, Genesse Scientific). Germ-free flies were generated by raising adult flies on autoclaved standard media supplemented with an antibiotic solution (100 μg/mL ampicillin, 100 μg/ml metronidazole, 40 μg/mL vancomycin dissolved in 50% ethanol and 100 μg/mL Neomycin dissolved in water) to eliminate the microbiome from adult flies. CR flies were raised on autoclaved standard cornmeal medium.

### Generation of mono-associated *Drosophila*

Virgin females were raised on selective medium for 5 days at 29° C. On day 5 of antibiotic treatment, a fly from each group to be mono/ploy-associated was homogenized in MRS broth and plated on MRS and GYC agar plates to ensure eradication of pre-existing microbes. Flies were starved in sterile empty vials for 2 hours prior to bacterial association. Lab isolate *Acetobacter pasteuranius* was grown in Mannitol broth at 29°C with shaking 2 days prior to association. Lab isolates *Lactobacillus brevis* and *Lactobacillus plantarum* were grown in MRS broth at 29°C 1 day prior to association. For mono-associations, the OD_600_ of bacteria liquid cultures was measured and then the culture was spun down and re-suspended in in 5% sucrose in PBS to final OD_600_ of 50 For poly-associations, bacterial cultures of *Ap, Lb*, and *Lp* were prepared to an OD_600_ of 50 in 5% sucrose in PBS as described above. The bacterial cultures were then mixed at a 1:1:1 ratio. For all bacterial associations, twelve flies/vial were associated with 1mL of bacterial suspension on autoclaved cotton plugs. Flies were fed the bacteria sucrose suspension for 16 hours at 29°C and then kept on autoclaved food for 5 days prior to infection. CR and GF flies were given mock associations of 1ml of 5% sucrose in PBS for 16 hours at 29°C. To ensure mono-association or GF conditions, respective flies were homogenized in MRS broth and plated on MRS or GYC agar plates.

### Colony forming units per fly

At indicated time points 25 flies per infection group were collected and placed into successive solutions of 20% bleach, distilled water, 70% ethanol, distilled water to surface sterilize and rinse flies respectively. These 25 flies were then randomly divided into groups of 5 and mechanically homogenized in LB broth. Fly homogenate was then diluted in serial dilutions in a 96 well plate and 10ul spots were then plated on either MRS agar (to select for *Lactobacillius*) GYC agar (to select for *Acetobacter*) and LB agar supplemented with 100 μg/ml streptomycin (to select for *Vibrio cholerae*).

### Competition Assays

Competition assays were performed as described in^13^. Briefly, *V. cholerae* V52 or V52Δ*vasK* were mixed with commensal bacteria at a 10:1 ratio and incubated on a pre-dried LB agar plate. A.fter a 2-h incubation at 37°C, bacteria were harvested, serially diluted and plated onto GYC or MRS agar to enumerate surviving commensals. Colony-forming units (CFUs) were counted the next day.

### Transmission Electron Microscopy

Flies were washed with 95% ethanol and dissected into PBS. Posterior midguts were immediately excised and placed into fixative (3% paraformaldehyde + 3% glutaraldehyde). Fixation preparation, contrasting sectioning, sectioning, and visualization were performed at the Faculty of Medicine and Dentistry Imaging Core at the University of Alberta. Midgut sections were visualized with Hitachi H-7650 transmission electron microscope at 60Kv in high contrast mode.

## Acknowledgements

*imd* mutant flies were provided by Bruno Lematire. The research was funded by Grants from the Canadian Institutes of Health Research to EF (MOP 77746) and SP (MOP 137106). We acknowledge the imaging and microscopy support from Dr. Stephen Ogg and Woo Jung Cho and the Faculty of Medicine and Dentistry core imaging service at the Cell Imaging Centre, University of Alberta.

